# Solving the SLoSS debate: Scale-dependent effects of habitat fragmentation on biodiversity loss

**DOI:** 10.1101/2024.08.27.609888

**Authors:** Monique de Jager, Edwin T. Pos

**Author notes:** Corresponding author:* Monique de Jager, Quantitative Biodiversity Dynamics, Department of Biology, Utrecht University, Netherlands. Statement of authorship:* MJ and EP designed the study. MJ created the model, analysed the simulation results, and wrote the first draft of the manuscript. EP provided critical feedback on model creation and data analysis, and contributed substantially to revisions of the manuscript. Data accessibility statement:* The source code of the model (R) and generated datasets are available at Zenodo (10.5281/zenodo.1337378).

## Abstract

Should what is left of nature be contained in a Single Large or Several Small (SLoSS) areas? This question of what would minimize the impact severity of habitat destruction on biodiversity loss is much debated, mainly because studies generally focus on different spatial and temporal scales. Using a semi-spatially explicit, (near-)neutral, individual-based model, we investigate the effects of fragmentation on biodiversity loss at two spatial (landscape-versus subcommunity level) and two temporal scales (static versus dynamic effects). Our results show that the role of spatial configuration of habitat destruction depends on when and at what scale we measure biodiversity loss. When considering the more realistic assumption that species differ in dispersal capacity, differences between spatial configurations are likely to be amplified. Our results indicate that the spatial configuration of habitat loss needs to be considered when evaluating the risks of further habitat destruction.

## Introduction

Habitat loss is a major issue that leads to a severe decline in biodiversity (Diamond 1989; Diffenbaugh and Field 2013; IPBES 2019). While the effects of habitat loss are commonly acknowledged, destruction of habitat remains an ongoing process (Ricketts 1999; Revenga et al. 2000; Haddad et al. 2015; Taubert et al. 2018). In general, habitat destruction follows a complex, fractal-like pattern, as a consequence of interactions between infrastructure and geographic characteristics of the landscape (Curtis 1956; Thorpe 1978; Gardner et al. 1987; Krummel et al. 1987; Sugihara and May 1990; With 1997; Hargrove et al. 2002; Ewers and Laurance 2006). It is clear that there will need to be some balance between habitat loss, restoration and the ever-growing human population; it is therefore vital that effects of the spatial configuration of habitat destruction on biodiversity, connectivity, and community dynamics is sufficiently considered in order to minimize the negative effects if we cannot avoid them.

The spatial configuration of habitat destruction may impact the severity of biodiversity loss; consensus on this subject, however, has not yet been reached. While some argue that fragmentation *per se* (i.e. the spatial configuration of a constant, i.e. fixed, amount of habitat, hereafter referred to as fragmentation; (Fahrig 2003; May et al. 2019) may have a negative effect on biodiversity (Hanski et al. 2013; Haddad et al. 2015), others reason that fragmentation’s effect on biodiversity might even be positive in some cases (Quinn and Harrison 1988; Tscharntke et al. 2002; Campos et al. 2012; Hanski et al. 2013; Keil et al. 2015; Fahrig 2017; Seibold et al. 2017). Yet another perspective proposes that the spatial configuration of habitat is not important at all (Fahrig 2003; Yaacobi et al. 2007; Fahrig 2013); the only thing that would matter is the amount of habitat available (the habitat-amount hypothesis, (Seibold et al. 2017)).

Three main drivers of these discrepancies in the effects of habitat fragmentation on biodiversity loss are mentioned in scientific literature. First, the effectiveness of the spatial configuration of habitat may depend on spatial species distributions (May et al. 2019): fragmentation may have a negative effect on biodiversity when species are regularly distributed, a positive effect when species are clustered, and no effect when species are randomly distributed. Second, we need to disentangle the effects of fragmentation on biodiversity that results in immediate species loss, often referred to as geometric or static fragmentation effects, and species loss after subcommunities have stabilized, generally referred to as demographic or dynamic fragmentation effects (Claudino et al. 2015). Most studies claiming positive effects on biodiversity concern immediate species loss of spatially aggregated species (e.g. (Quinn and Harrison 1988; Seibold et al. 2017; May et al. 2019)); as more distant subcommunities generally have quite a different species composition, random habitat destruction results in the lowest immediate species loss at the landscape scale (Claudino et al. 2015; Chisholm et al. 2018). Third and last, consensus on effects of fragmentation has not been reached as studies drawing different conclusions generally also focus on different spatial scales, confounding any generalization of observations. As a result of these different drivers and the lack of generalization, the effects of habitat fragmentation on subcommunity dynamics and connectivity remains quite elusive.

The above is summarized in the ongoing SLOSS debate – whether Single Large Or Several Small habitat fragments are better for the conservation of biodiversity – emphasizing that a better understanding is vital, being it on the effect of scale on which results are measured, or disentangling static and dynamics fragmentation effects (May et al. 2019). Here, we investigate the effects of both drivers of disagreement between studies: static versus dynamic and landscape versus subcommunity-level effects of fragmentation on biodiversity loss. Additionally, we examine the effect of differences in species’ dispersal capacity, as it is known that dispersal capacity has a substantial effect on community dynamics and connectivity and hence presumably also interacts with the consequences of habitat fragmentation (de Jager and Soons 2023). We created a semi-spatially explicit, (near-)neutral, individual-based model in which destruction of habitable areas is simulated in three types of spatial configuration: (i) spatially clustered, (ii) fractal, and (iii) random habitat destruction.

We hypothesize that, while random habitat destruction may result in a smaller immediate (static) species loss at the landscape scale, it will result in a steeper decline in species numbers with increasing habitat loss, compared to fractal or clustered habitat destruction, after subcommunities have stabilized (dynamic, subcommunity-level). Especially in random configurations of habitat destruction, the dynamic fragmentation effects caused by, for example, Allee effects (Courchamp et al. 2008; Swift and Hannon 2010), demographic stochasticity (MacArthur and Wilson 1967; Lande 1993), and reduced migration (Hanski 1999; Hanski et al. 2013) are expected to result in severe biodiversity loss. We furthermore hypothesize that species composition changes from long-distance dispersers in unfragmented environments to higher occurrences of locally dispersing species in environments with increased habitat loss. This change in species composition may subsequently impact community dynamics, connectivity, and further biodiversity loss.

## Methods

We simulated subcommunity dynamics in a 2-dimensional, semi-spatial, near-neutral, individual-based model. The environment consists of 20 x 20 cells, each cell having a subcommunity of 1,000 individuals. Initially, all cells are habitable (i.e. there is no fragmentation) and contain a subcommunity. Each subcommunity is populated with the highest diversity theoretically possible during an initialization phase and is subsequently allowed to stabilize, after which we run the simulations with habitat loss. We simulated 20 different levels of habitat loss, from 0 to 95% habitat destruction, at three fragmentation configurations: (i) clustered, (ii) fractal, and (iii) random habitat destruction. Each simulation was run 10 times to account for the stochastic nature of the model. All model simulations and analyses of the resulting data were performed in R version 4.3.1 using the following packages: ggplot2, future, meteR, and benthos (van Loon et al. 2015; Wickham 2016; Rominger and Merow 2017; Bengtsson 2021).

### Creating initial communities

Subcommunities were initialized in the modelled landscape. We created a 2-dimensional lattice of 20 x 20 cells with continuous boundaries (i.e. a torus), where each cell contained a subcommunity consisting of 1,000 individuals. Each subcommunity started with one individual per species of species S_1_ to S_1,000_ to maximize initial diversity. Each timestep, every individual was able to disperse within its own or to a neighboring subcommunity and randomly replace another individual. The Euclidean distance between cells determined the probability that an individual from subcommunity *i* could replace an individual in subcommunity *j* (Fig. S1). To limit matrix sizes and thereby increase computation speed, dispersal was limited to the closest 121 cells (within the ranges -5 ≤ *Δx_ij_* ≤ 5 and -5 ≤ *Δy_ij_* ≤ 5, where *Δx_ij_* and *Δy_ij_* are the distances (in numbers of cells) in *x* and *y* directions between the focal subcommunity *i* and its neighbor *j*). For each of these 121 cells surrounding (and including) a focal subcommunity, the probability to disperse here was calculated using a 2-dimensional exponential probability density function:

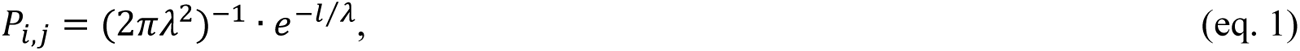

where *λ* is the scaling exponent and *l* is the Euclidian distance between subcommunities *i* and *j*. Other probability density functions can also be used, to describe different dispersal kernels. We ran two different simulation blocks, one in which all species dispersed with λ = 0.8, hereafter called same-dispersal simulations, and one in which species differed in their dispersal capacity (0 < λ < 1), hereafter called different-dispersal simulations. For the same-dispersal simulations, λ = 0.8 was chosen, as this was the median dispersal capacity of the individuals in the different-dispersal simulations within a pristine environment.

Every individual in each subcommunity was randomly assigned a subcommunity to disperse to, based on its dispersal kernel (weighted random selection on *P_i,j_*). The individual to replace from that subcommunity was randomly selected as well (without any differences in weights depending on for example the species). Because such random replacements may overlap (i.e. an individual was replaced twice in the same iteration) and R always uses the last value as the resulting replacement, we randomly ordered the replacements. Replacement of the individuals in the subcommunities was carried out for 10,000 iterations.

### Fragmenting the environment

We simulated increasing habitat loss in steps of 5% habitat destruction (resulting in 20 sequential environments with different fragmentation levels), and with 3 different levels of clustering of habitat destruction. For each landscape, the first non-habitat cell was randomly selected from all cells. Subsequently, while the number of non-habitat cells created was smaller than the desired number of non-habitat cells (which depends on the simulated level of habitat loss), non-habitat cells were sequentially selected by means of weighted random selection, where weights of cells were attributable to their distance from destructed, non-habitat cells. Weights (*W_i_*) were calculated from distances using a 2-D pareto distribution:

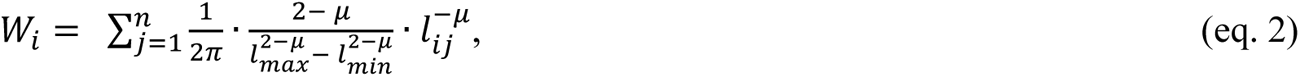

where *l_ij_* is the Euclidian distance between subcommunity *i* and non-habitat cell *j*, *n* is the number of non-habitat cells, *l_min_* and *l_max_* are the minimum and maximum distances to consider (*l_min_* = 1, *l_max_* = 50), and *µ* is the scaling exponent (*µ* > 1). When *µ* is small, the non-habitat cells are distributed more homogeneously; non-habitat cells become more clustered with increasing *µ*. The different spatial configurations of habitat destruction were created using *µ* = 1 for random, *µ* = 3 for fractal, and *µ* = 5 for clustered habitat destruction (Fig. 1).

**Figure 1:**
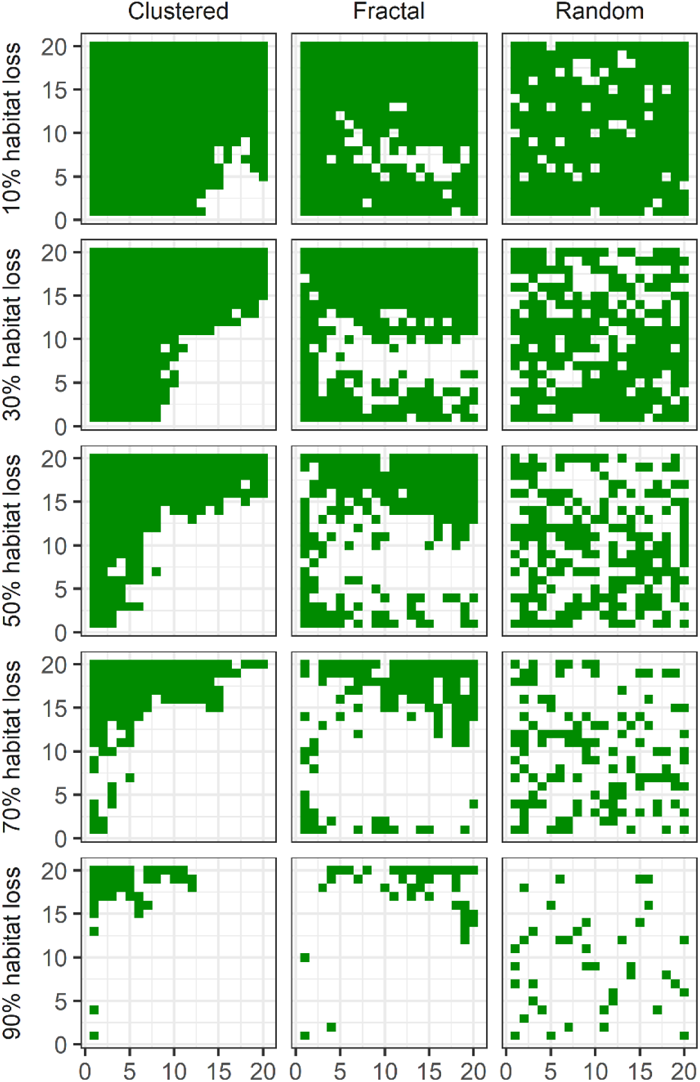
Some examples of the spatial configuration of remaining subcommunities (green patches) after clustered, fractal, or random habitat destruction has occurred.

The spatial configuration of the environments with clustered, fractal, and random habitat destruction differed substantially from each other (Fig. S2). Under random habitat destruction, Moran’s I, an indicator of spatial autocorrelation, was close to zero, the average patch size decreased rapidly with habitat loss, and the number of patches showed a large peak at 75% habitat loss. In contrast, under clustered habitat destruction, spatial autocorrelation was high, mean patch size decreased less rapidly with habitat loss, and the number of patches remained much more constant across the different levels of habitat loss.

### Community dynamics

Simulations with different amounts of habitat loss started with the communities that were recorded at the end of the initialization round (e.g. after 10,000 iterations in a pristine environment). During a simulation, all individuals iteratively reproduced and replaced other individuals, as described above in the section *Creating initial communities*. With a probability of 0.0003 (Pos et al. 2017), individuals coming from a larger (hypothetical) metacommunity settled in the subcommunities, where the individual’s species (*S_1_* to *S_1,000_*) was selected using weighted random sampling that depended on the species’ dispersal capacity.

### Data analysis

For each simulation, we recorded – per subcommunity – (i) the number of species, (ii) the fraction of individuals originating from ancestors that inhabited the current patch 50 timesteps earlier (t_-50_), and (iii) the fit of the prediction of Maximum Entropy Theory in Ecology (METE) to the species abundance distribution (SAD) (Harte and Newman 2014). With METE, a large number of different community relations and distributions can be estimated, often more accurately than by any other method (Harte et al. 2008; Rominger and Merow 2017). Using only the average number of individuals per species and the total number of individuals as input (called constraints), METE can predict the species abundance distribution (SAD) very accurately for undisturbed, steady-state ecosystems (Harte et al. 2008). Yet, for disturbed systems, METE estimations deviates from observed distributions (Newman et al. 2020). We propose that, as METE estimates deviate further in more disturbed systems, we can use this deviation as a measure of ecosystem disturbance. We estimated SAD using the *sad* function from the package *meteR* (Rominger and Merow 2017) on the sampled data. For the given number of species present, we generated 1,000 SAD’s and calculated the average abundance per rank. Using this average estimated SAD and the actual simulated SAD, we calculated the goodness of fit (GOF_METE_):

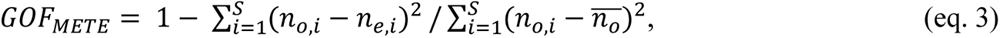

where *S* is the number of species present in the local community, *n_0,i_* and *n_e,i_* are, respectively, the number of individuals observed and expected (by the METE estimation) for rank *i*, and 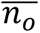 is the average number of individuals per species. Values of *GOF_METE_* close to 1 indicate that the prediction by METE of the SAD is highly comparable and, hence, the subcommunity appears to be a in state that would adhere to what is expected based on the macroscopic constraints. We furthermore calculated Bray-Curtis dissimilarity (using the bray_curtis function from the benthos R-package on the counts per species at each site), after each simulation round, between 20 randomly selected subcommunities.

## Results

### Landscape-scale species-area relations

On a landscape-scale, static (geometric) species loss due to habitat loss was less in environments with random habitat destruction than in those with clustered destruction of habitat cells (Fig. 2, top panels). Fewer species were lost in same-dispersal simulations than in different-dispersal simulations. In different-dispersal simulations, all three spatial configurations resulted in species-area curves less steep than that of the backwards SAR (the species-area relation observed in the pristine, unfragmented environment). In contrast to immediate species loss, dynamic (demographic) species loss was much larger than what was predicted by the backwards SAR, resulting in species-area curves declining faster than that derived after 10,000 iterations of community dynamics in the pristine landscape (Fig. 2, bottom panels). For same-dispersal simulations, random habitat destruction still resulted in the lowest species loss with increasing habitat loss (e.g. decreasing area size); yet, in different-dispersal simulations, random habitat destruction at intermediate levels of habitat loss resulted in the highest amount of species loss. In all cases, increasing habitat loss resulted in a decreased number of species on the landscape-scale.

**Figure 2:**
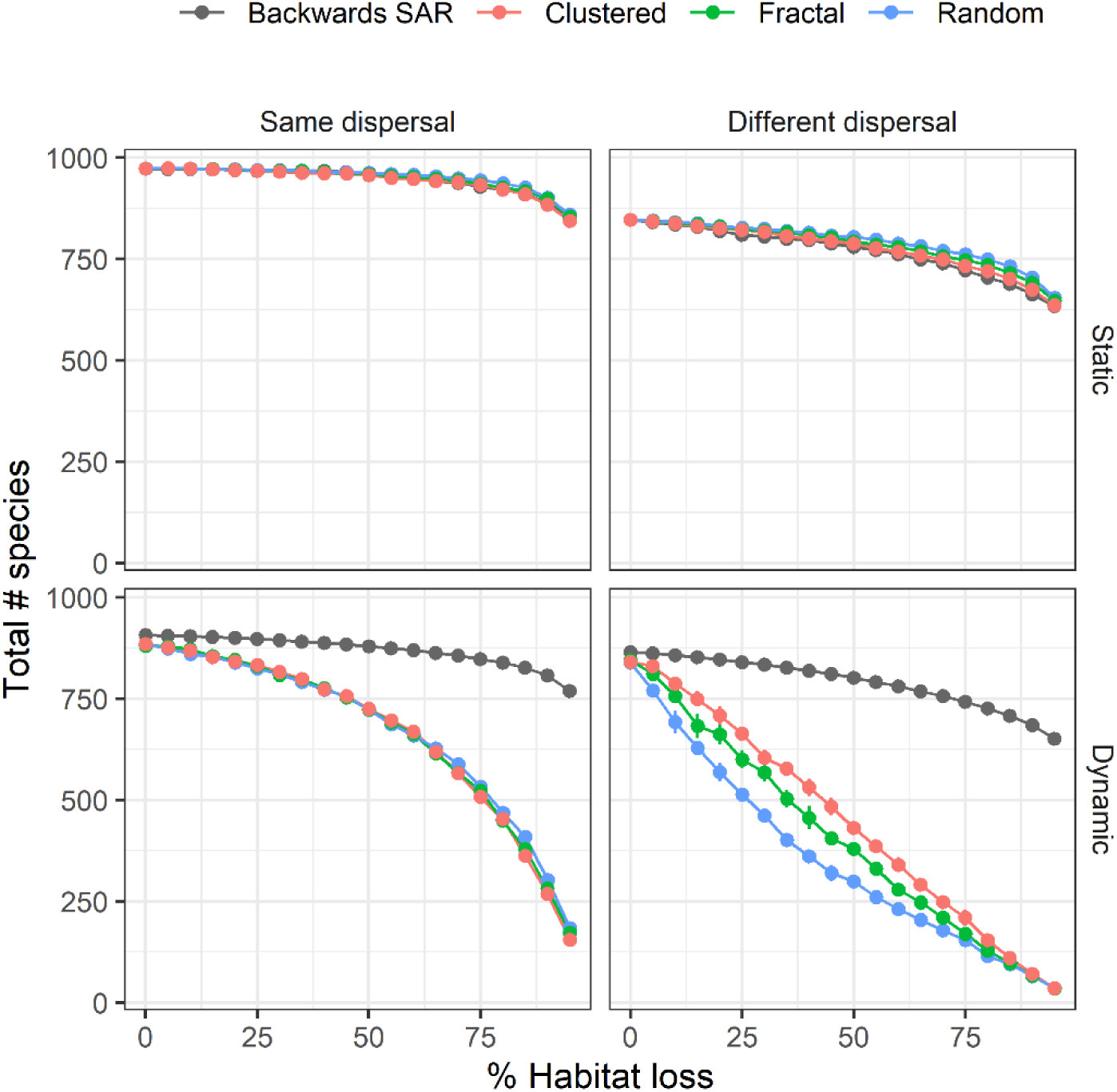
Total number of species present immediately after habitat loss (Static, top panels) and after community dynamics have stabilized (Dynamic, bottom panels) in the simulated environment per percentage habitat loss, for different spatial configurations of habitat destruction (clustered = red, fractal = green, and random habitat destruction = blue) and in simulations in which species have the same (left panels) or different dispersal strategies (right panels). The grey lines indicate the species-area relations in the pristine, unfragmented environments. Error bars indicate the 95% confidence interval.

### Subcommunity-scale effects of habitat loss and fragmentation

*Same vs. different dispersal simulations.* At the level of the subcommunity, we did, however, observe major differences in dynamic effects of habitat loss and fragmentation. First, there were differences between simulations with same versus different dispersal strategies. In different-dispersal simulations, a shift occurred from long-distance dispersers being more prevalent in continuous habitat, towards local dispersing species when habitat destruction was introduced (Fig. 3). This shift in dispersal capacity had a clear impact on subcommunity connectivity and dynamics. In same-dispersal simulations (Fig. 4, top-left panel), the number of species per subcommunity decreased at a much slower rate than in different-dispersal simulations (Fig. 4, top-right panel). The percentage of t_-50_ ancestors from outside of the focal subcommunity remained high for most habitat loss levels in case of clustered and fractal habitat destruction in same-dispersal simulations (Fig. 4, center-left panel), while this percentage decreased rapidly with habitat loss in different-dispersal simulations (Fig. 4, center-right panel). Furthermore, METE’s fit to SAD was substantially better at high levels of habitat loss in same-dispersal (Fig. 4, bottom-left panel) than in different-dispersal simulations (Fig. 4, bottom-right panel).

**Figure 3:**
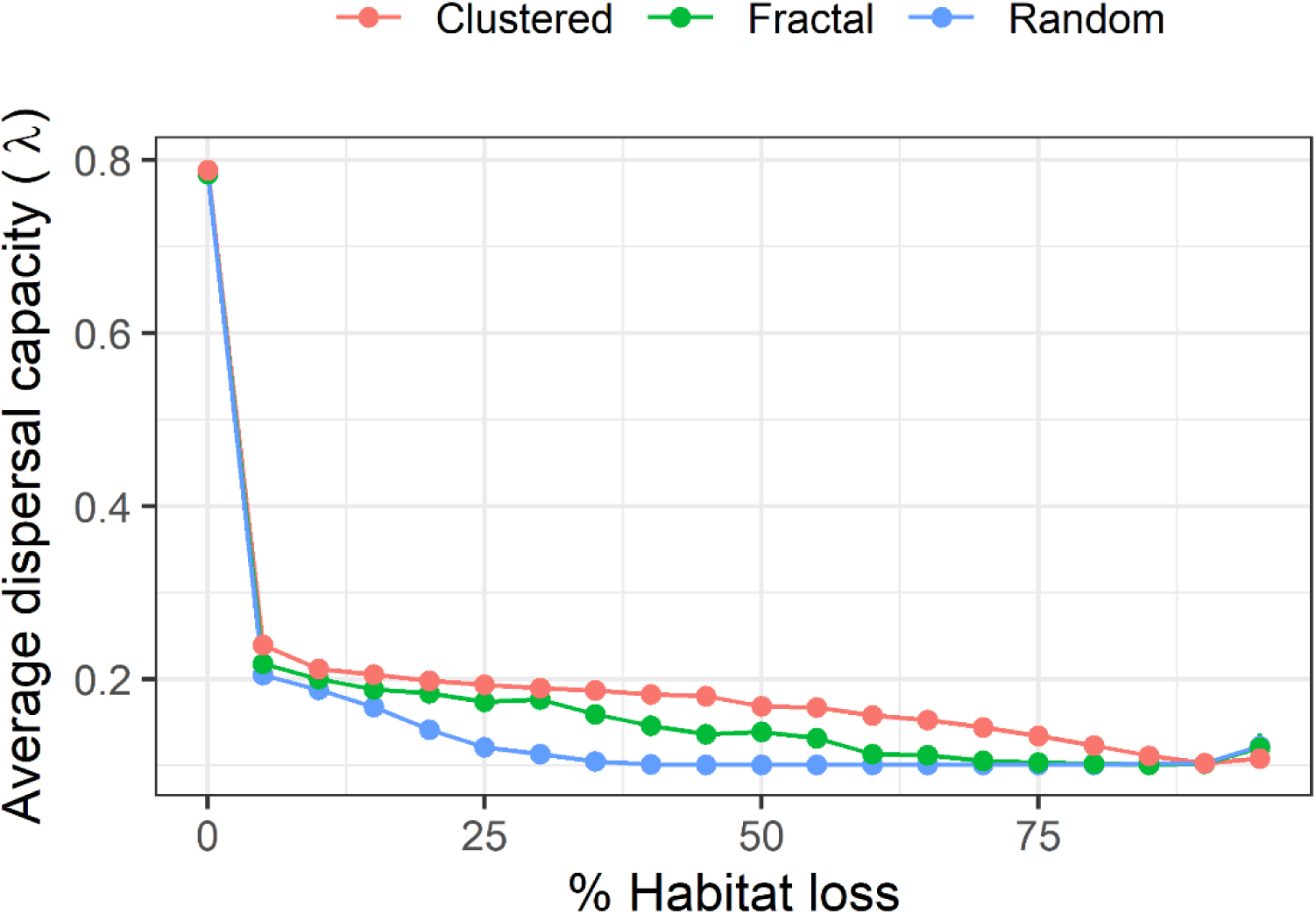
The average dispersal capacity (λ) is shown per percentage habitat loss and for the three different spatial configurations of habitat destruction (colors: clustered = red, fractal = green, and random habitat destruction = blue). Error bars indicate the 95% confidence interval.

**Figure 4:**
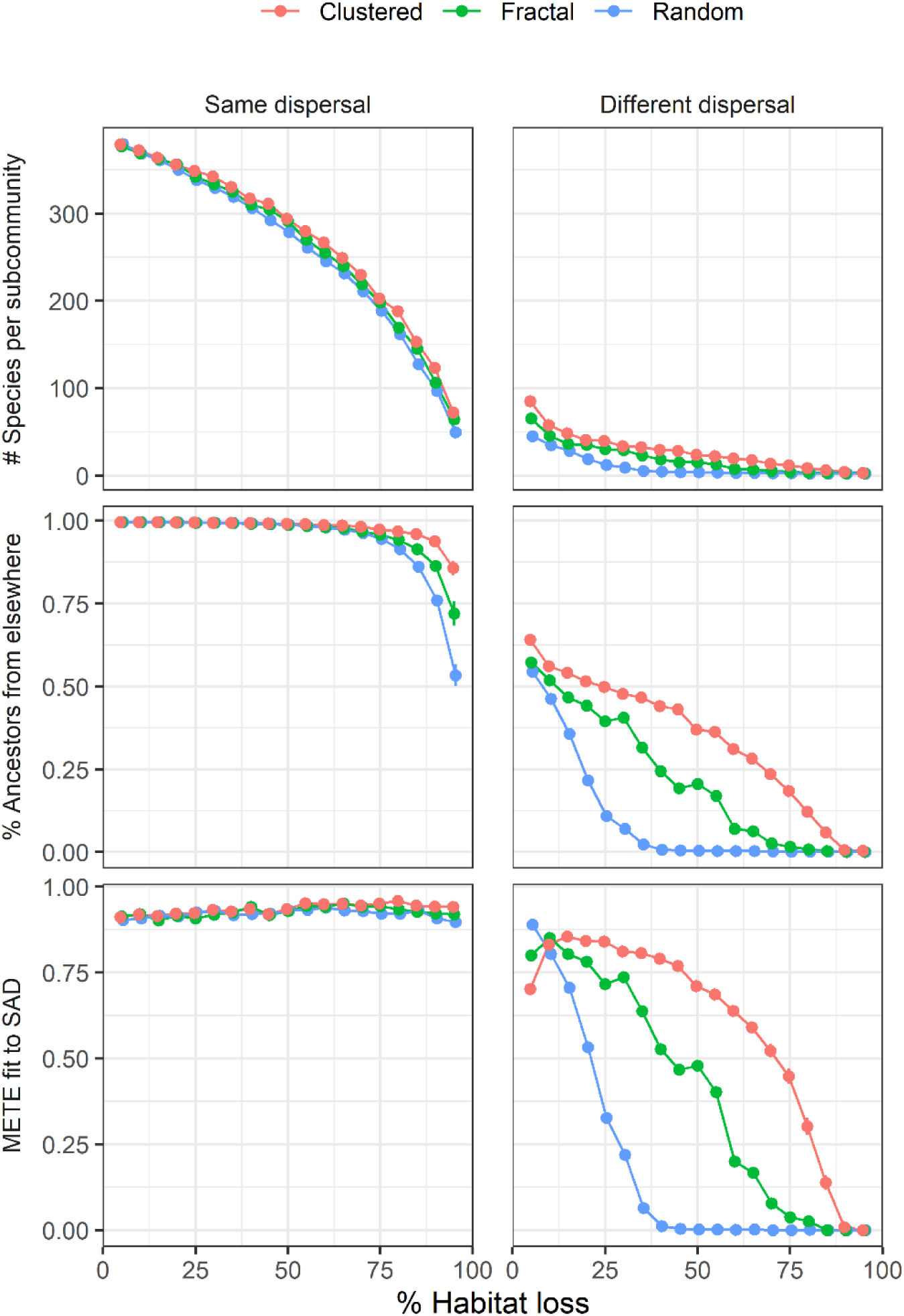
The average number of species per subcommunity (top panels), the average percentage of t_-50_ ancestors from another subcommunity per subcommunity (center panels), and the average METE fit to SAD per subcommunity (bottom panels), for the different combinations of the percentage of habitat loss (x-axis) and the three different spatial configurations (colors: clustered = red, fractal = green, and random habitat destruction = blue), when all species are using the same (long-distance) dispersal strategy (left panels), or when species differ in their dispersal strategy (right panels). Error bars indicate the 95% confidence interval.

*Effects spatial configuration of habitat destruction.* Second, the spatial configuration of habitat destruction affected community dynamics and connectivity in both same-dispersal and different-dispersal simulations (Fig. 4, columns). Random habitat destruction had a larger deleterious effect on subcommunity connectivity and dynamics than clustered or fractal distributions of habitat destruction. With a random distribution of habitat destruction, the number of species per subcommunity decreased more quickly with increasing habitat loss (Fig. 4, top panels). The percentage of t_-50_ ancestors from outside of the current subcommunity decreased more rapidly with increasing habitat loss in environments with random destruction as well (Fig. 4, center panels). Furthermore, the METE fit to SAD per subcommunity quickly transitioned from a good fit towards zero with increasing habitat loss in environments with random habitat destruction in different-dispersal simulations (Fig. 4, bottom panels).

Bray-Curtis dissimilarity in terms of species composition between subcommunity pairs was also affected by habitat loss, spatial configuration, and variation in dispersal strategies (Fig. 5). In same-dispersal simulations, subcommunities became more similar with increasing habitat loss when habitat destruction was clustered. In contrast, dissimilarity increased with habitat loss in same-dispersal simulations with random habitat destruction, as well as in all different-dispersal simulations, regardless of their spatial configuration. In different-dispersal simulations, we observed another effect of spatial configuration on the strength of the impact of habitat loss; with random habitat destruction, dissimilarity increased more rapidly than with fractal or clustered habitat destruction.

**Figure 5:**
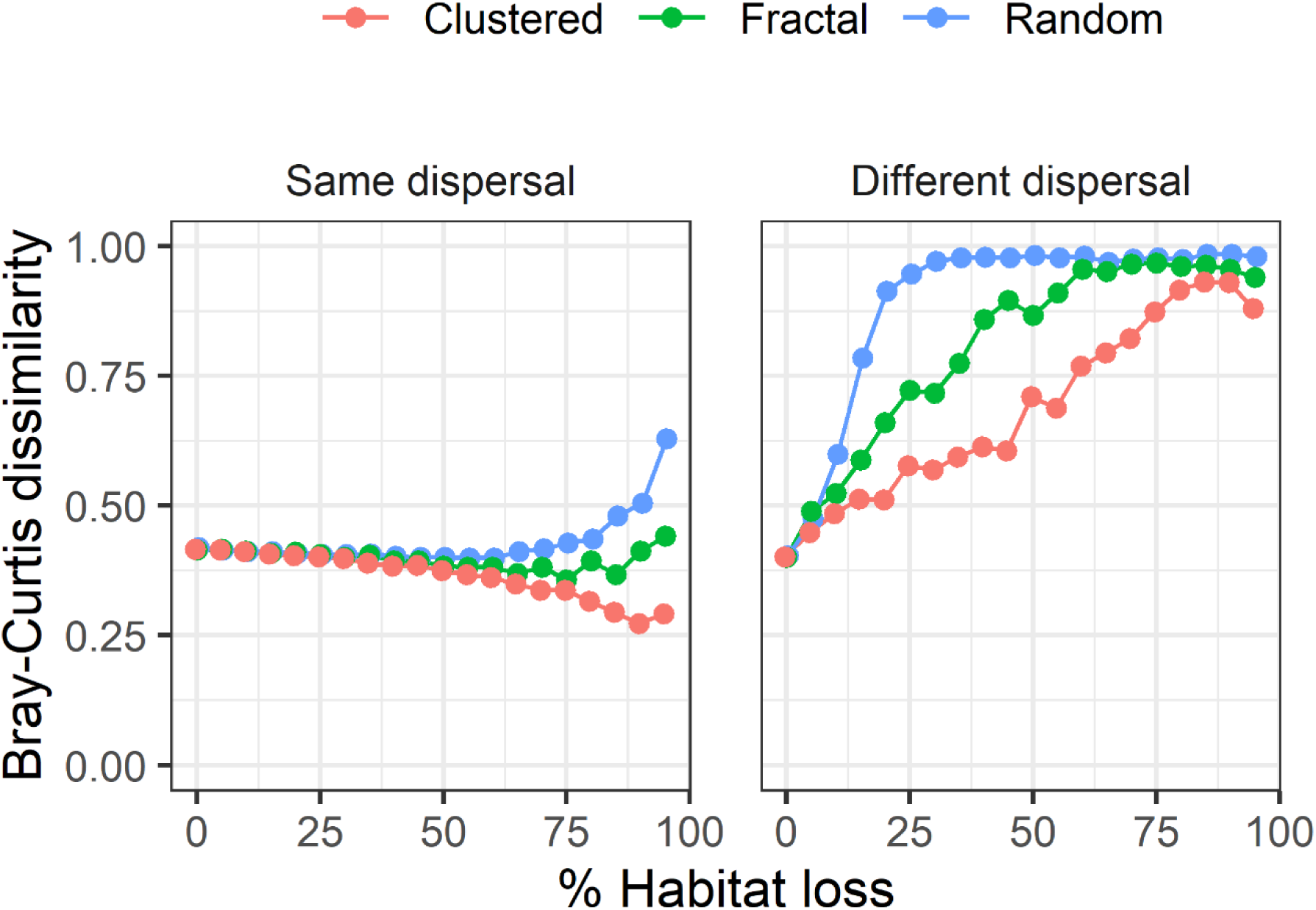
The average Bray-Curtis dissimilarity for different levels of habitat loss (x-axes), spatial configuration of habitat destruction (colors: clustered = red, fractal = green, and random habitat destruction = blue), in same- and different-dispersal simulations (panels). Error bars indicate the 95% confidence interval.

## Discussion

While it is a truth universally acknowledged that habitat loss generally also results in biodiversity loss, the role of the spatial configuration of habitat or habitat destruction, however, has been often debated. Here, we show that the role of spatial configuration of habitat destruction, or habitat fragmentation, depends on when and at what scale we measure biodiversity loss. Similar to other studies (Campos et al. 2012; Claudino et al. 2015; Keil et al. 2015; May et al. 2019), our results show that static biodiversity loss (i.e. loss of species immediately after habitat areas have been destructed), measured at the spatial scale of the entire environment, is smallest in case of random habitat destruction. However, after stabilization of subcommunity dynamics, this advantage of a random configuration over fractal and clustered habitat destruction disappears. Moreover, at the much smaller scale of single subcommunities, biodiversity loss caused by habitat loss is worse in the case of random habitat destruction than in case of less habitat fragmentation.

Whereas species-area curves were less steep than the backwards SAR in case of static fragmentation effects, they were substantially steeper after communities had stabilized. Many species that have evolved in continuous, pristine landscapes are maladapted to life in small habitat patches, and are therefore likely to go extinct locally (Simberloff 1992). The extinction debt – i.e. the future extinction of species due to the habitat destruction that has taken place – is often assumed to settle towards a zero deviation from the backwards SAR after dynamic fragmentation effects have stabilized the community (Claudino et al. 2015), Yet, our results show that species numbers can drop far below what is expected from backwards SAR when habitat is lost. This is a serious reason for concern, advocating once more for the discontinuation of habitat destruction to stop further increase in biodiversity loss.

Dispersal plays a crucial role in connecting subcommunities. With our model, we ran two types of simulations, one in which all species had an equal dispersal propensity (as this is the default approach to model community dynamics in a neutral model (Hubbell 2001; Chisholm et al. 2018), and one in which species differed in their dispersal capacity (i.e. a near-neutral approach). In the more realistic setting of species differing in their dispersal capacity, species composition changed as a result of habitat loss and fragmentation, from long-distance to short-distance dispersers becoming more prevalent in the subcommunities. Due to this shift, connectivity between subcommunities worsened, subcommunities became more dissimilar, and species were lost at a fast rate.

The absence of substantial differences in dynamic biodiversity loss at the landscape-scale can be attributed to the differences in species numbers per subcommunity and subcommunity dissimilarity between the different spatial configurations cancelling each other out. Subcommunities in simulations with clustered habitat destruction have a larger number of species per subcommunity and are also more similar in species composition than those in simulations with random habitat destruction. These species are thus often the same species as in other subcommunities, resulting in a lower number of species at the landscape-scale than when subcommunities would have been less similar in species composition. In contrast, subcommunities in simulations with random habitat destruction have low species numbers but high dissimilarity between subcommunities. As these few species are often different from the species in other subcommunities, the number of species at the landscape-scale is higher than expected if subcommunities were to be more similar.

As models are simplified representations of reality, they have their limitations. Some of our model’s limitations include the absence of spatial heterogeneity in habitat quality and habitat types, the absence of interactions between species, such as predator-prey interactions, plant-pollinator, and plant-seed dispersers interactions, and the absence of niche differentiation between species. Species in our model are unaffected by edges, other than them dispersing into the uninhabitable matrix. Time between habitat destruction events, as well as the magnitude of habitat loss per event, presumably will also considerably affect community dynamics and biodiversity loss (Claudino et al. 2015). Further investigation is thus needed to explore these effects. Furthermore, we did not explicitly examine the effect of the spatial distribution of species (which is one of the three aspects assumed to impact the effect of habitat fragmentation on biodiversity). In our simulations, species can differ in their dispersal strategy and hence in their degree of spatial clustering. Species with a high dispersal capacity are more randomly distributed than those with short-distance dispersal, which are more clustered. The clustered, short-distance dispersing species will, at first, have an advantage over the more randomly distributed, long-distance dispersing species when habitat is lost, especially under random habitat destruction (May et al. 2019). This process may also explain the increase in short-distance dispersers when habitat is lost.

The model presented here can be a useful tool for future research. With this model, the effects of time between habitat loss events, as well as their size, can be investigated more thoroughly. Model outcomes can also be compared with those of experimental studies. While most empirical studies on habitat loss and fragmentation lack information on community composition in the original, unfragmented landscapes, some experimental studies do; their respective timeframes, however, are still quite short. By linking model results to those from experimental studies, we can predict species loss in the future, using a dynamic framework based on time series. In addition, changes in METE fit to SAD after habitat destruction and during the process of stabilization of fragmented subcommunities could be explored further, as this might be a good indicator of community stability.

Concluding, the best solution would obviously be to have no habitat destruction at all, yet sometimes this either is unavoidable or compromises must be made given the many conflicts of interests between a growing population and a limited amount of space available. While habitat loss is always detrimental to biodiversity, our results suggest that fewer species are lost at the subcommunity level when habitat destruction is more clustered. In turn, this suggests that the damage done to biodiversity can at least somewhat be lessened by considering the spatial configuration of habitat destruction and we urge the community to take this into account, now and in the future.

---

**Text box.**
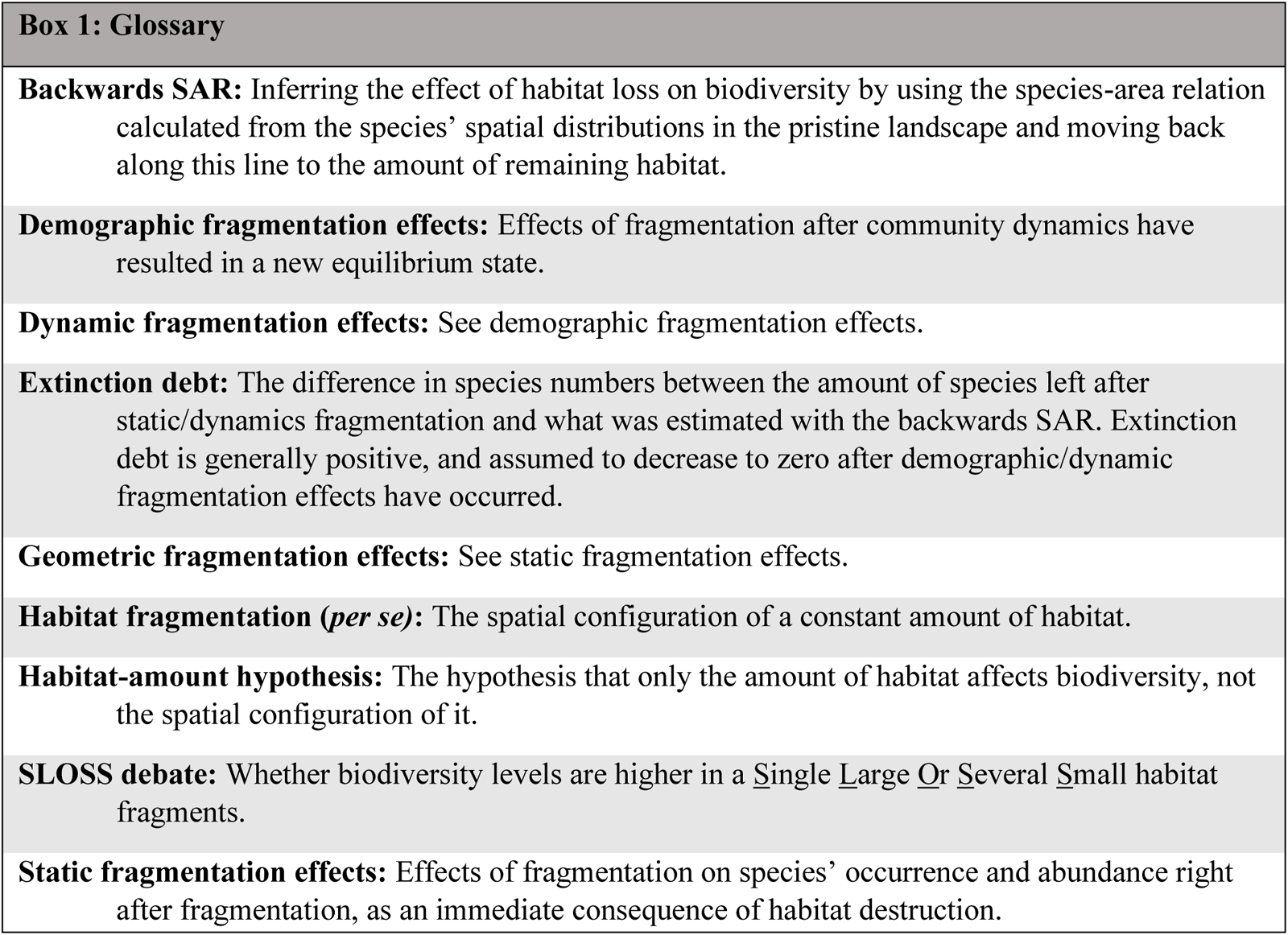

