## Supplementary Figures for "Solving the SLoSS debate: Scale-dependent effects of habitat fragmentation on biodiversity loss"

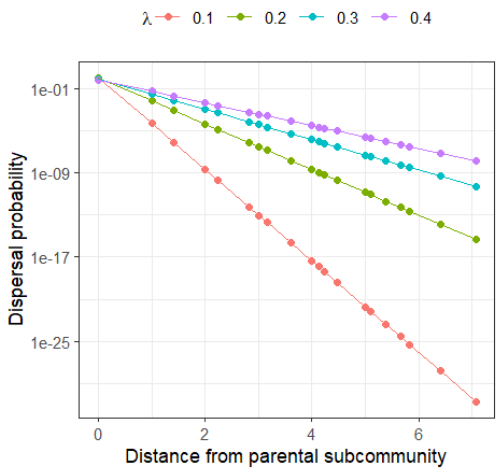


***Figure S1:*** *Weighted random sampling of the subcommunity to disperse to was based on the dispersal probability, which in turn depended on the distance from the parental subcommunity and the dispersal exponent λ.*

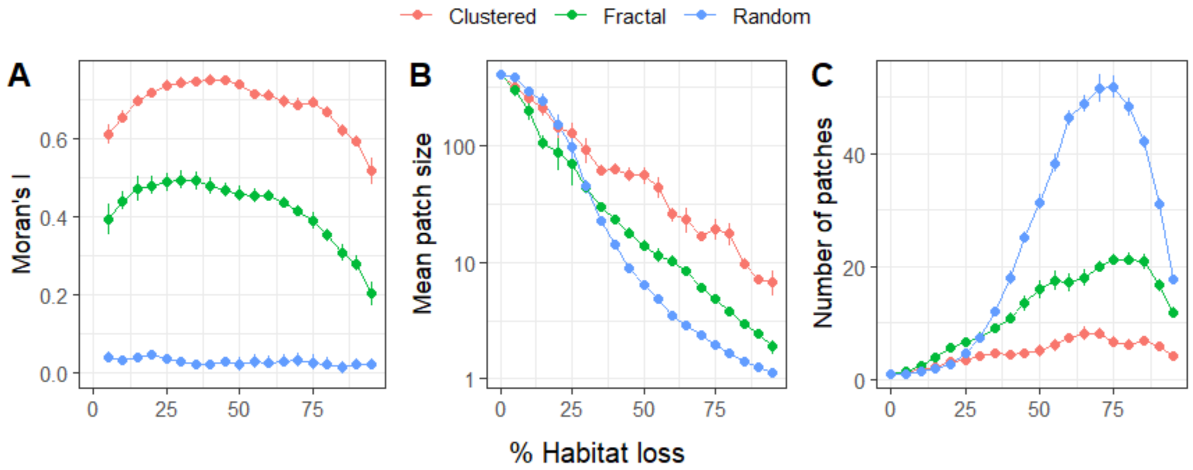


***Figure S2:*** *(**A**) Moran's I (an indicator for spatial autocorrelation), (**B**) mean patch size, and (**C**) the number of patches per percentage habitat loss and spatial configuration (clustered (red), fractal (green), or random (blue) habitat destruction).*
